# Immobilisation of Lipophilic and Amphiphilic Biomarker on Hydrophobic Microbeads

**DOI:** 10.1101/2023.01.10.523433

**Authors:** Franziska Dinter, Thomas Thiehle, Uwe Schedler, Werner Lehmann, Peter Schierack, Stefan Rödiger

**Affiliations:** Brandenburg University of Technology Cottbus-Senftenberg, Universitätsplatz 1, 01968 Senftenberg, Germany; PolyAn GmbH, Schkopauer Ring 6, 12681 Berlin, Germany; Freie Universität Berlin, Takustraße 3, 14195 Berlin, Germany; attomol GmbH, Schulweg 6, 03205 Bronkow, Germany; Faculty of Health Sciences, joint Faculty of the Brandenburg University of Technology Cottbus – Senftenberg, the Brandenburg Medical School Theodor Fontane and the University of Potsdam, Berlin, Germany; CreativeOpenLab (COLab), Siemens-Halske-Ring 2, 03044 Cottbus, Germany

**Keywords:** microbeads, hydrophobic, phospholipids, amphiphilic biomarker

## Abstract

**Background:** Lipids and amphiphilic molecules are ubiquitous and play a central role in cell signalling, cell membrane structure, and lipid transport in the human body. However, they also appear in many diseases such as atherosclerosis, cardiovascular diseases, infections, inflammatory diseases, cancer, and autoimmune diseases. Thus, it is necessary to have detection systems for lipids and amphiphilic molecules. Microbeads can be one of these systems for the simultaneous detection of different lipophilic biomarkers.

**Methods:** Based on the fundamentals of microbead development, novel hydrophobic microbeads were produced. These not only have a hydrophobic surface, but are also fluorescently encoded and organic solvent resistant. The challenge after the development of the hydrophobic microbeads was to immobilise the amphiphilic molecules, in this study phospholipids, on the microbead surface in an oriented direction. After successful immobilisation of the biomarkers, a suitable antibody based detection assay was established.

**Results:** By passive adsorption, the phospholipids cardiolipin, phosphatidylethanolamine and phosphatidylcholine could be bound to the microbead surface. With the application of the enzymes phospholipase A2 and phospholipase C, the directional binding of the phospholipids to the microbead surface was demonstrated. The detection of directional binding indicated the need for the hydrophobic surface. Microbeads with no hydrophobic surface bound the phospholipids non-directionally (with the hydrophilic head) and were thus no longer reactively accessible for detection.

**Conclusion:** With the newly developed hydrophobic, dual coded and solvent stable microbeads it is possible to bind amphiphilic biomolecules directionally onto the microbead surfaces.

## Introduction

Lipids are ubiquitous in the human body and play a role in cell signalling, cell membrane structure, and lipid transport (Burdge and Calder, 2015). The lipids themselves or changes in the lipids are associated with diseases such as atherosclerosis (Solnica et al., 2020), cardiovascular diseases (Dinter et al., 2019; Plüddemann et al., 2012), infections and inflammatory diseases (Filippas-Ntekouan et al., 2017), Alzheimer’s disease and diabetes (Wang et al., 2019), as well as with cancer (Ahmadpour et al., 2020; Ye et al., 2021; Zou et al., 2021), autoimmune diseases (Cervera, 2017; Jizzini et al., 2020; Sebastiani et al., 2016), COVID-19 (Das, 2020; Jizzini et al., 2020), and neurological diseases (Falabella et al., 2021). In cancer, tumour cells have a high demand for phospholipids to build the cell membrane due to their rapid proliferation (Ye et al., 2021). In autoimmune diseases such as antiphospholipid syndrome, antiphospholipid autoantibodies play a role, e.g. against cardiolipin, ß-2GP1 or phosphatidylethanolamine. This may lead to arterial or venous thrombosis, thrombocytopenia, or pregnancy morbidity (Cervera, 2017; Faricelli et al., 2008; Galli et al., 2003; Sebastiani et al., 2016). Conventional methods for the detection of lipids are agarose gel electrophoresis (Kuromori et al., 2006), chromatography including mass spectrometry and liquid chromatography (Fernandes et al., 2020; Hidaka et al., 2007; Lange et al., 2019; Wang et al., 2019), shot-gun lipidomics (Postle, 2012), strip tests (Huang et al., 2020; Wang and Hu, 2020), ELISA (Dominiczak and Caslake, 2011; Giannakopoulos et al., 2009; Loizou et al., 1985), and various types of biosensors (Dimitrijevs and Arsenyan, 2021; Lu et al., 2017). Methods like lipid-ELISA are costly and often cannot identify several lipids simultaneously. However, the latter is important as sample material is limited (Tighe et al., 2015). Since their role in the diagnosis of diseases is becoming increasingly relevant, it is necessary to identify lipid and amphiphilic molecules quickly and with as little sample material as possible (Dinter et al., 2019). Here the use of microbeads lends itself, as these can be used for rapid and patient material saving analysis by allowing several biomarkers/parameters to be analysed simultaneously (Dinter et al., 2019; Rödiger et al., 2012b). Lipids are predominantly hydrophobic and soluble in organic solvents and therefore do not behave in the same way as others, e.g. hydrophilic biomarkers like proteins or nucleic acids (Burdge and Calder, 2015; Garcia et al., 2017). There is no possibility to bind the lipids to the surface of carboxylated microbeads via functional groups. Immobilisation of lipophilic and amphiphilic molecules requires high solvent stability of the microbeads. Fluorescently and size encoded, solvent stable microbeads with a hydrophobic surface do not yet exist. In this study, following the development of the hydrophobic microbeads, a method was established to immobilise the amphiphilic phospholipids cardiolipin, phosphatidylethanolamine and phosphatidylcholine to the microbead surface in a directed manner (binding with the hydrophobic tail).

## Material and Methods

### Material

#### Phospholipids

cardiolipin - Cy5, phos-phatidylethanolamine - Cy5, phosphatidylcholine - Cy5, were purchased from Avanti Polar Lipids, INC, Birmingham, Alabama. Phospholipases were purchased from Sigma Aldrich, St. Louis, USA. Anti human IgG antibody with Alexa-Fluor 647 was purchased from Dianova, Hamburg, Germany. Chemicals acetone, methanol and chloroform were purchased from Carl Roth GmbH+ Co.KG, Karlsruhe, Germany. The 200 μL tubes were purchased from Brand^®^, Wertheim Germany. Microbeads were provided by PolyAn GmbH, Berlin, Germany. Devices used were: thermo shaker (ThermoMixer C) from Eppendorf, Hamburg, Germany; centrifuge (Minispin plus), Eppendorf, Hamburg, Germany; contact angle metre (OCA 30) from dataphysics, Filderstadt, Germany.

### Microbeads

We used six different poly(methylmethacrylat) (PMMA) microbead populations. Microbeads were coded with two fluorescence dyes (Rhodamine 6G and Coumarin 6) and different surface modifications (Table 1). MB1-MB4 are hydrophobic and MB5 and MB6 are hydrophilic, due to their surface modifications.

**Table 1.**
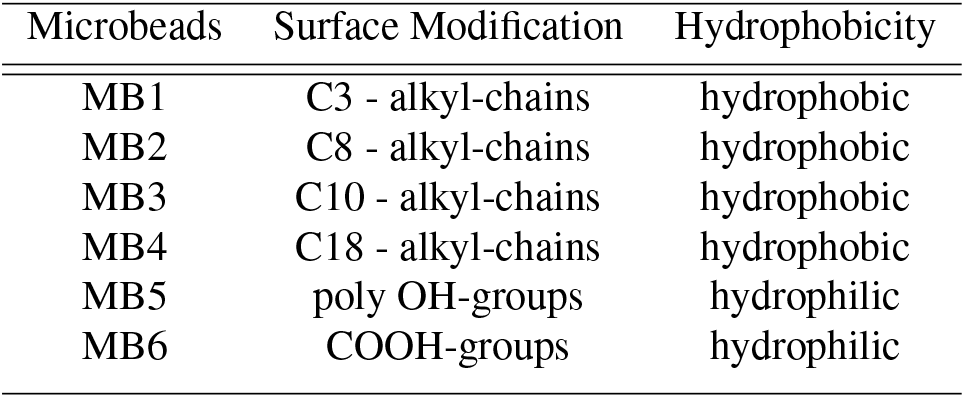
Microbead Populations and their characteristics.

### Solvent stability of carboxylated and hydrophobic microbeads

To test the stability of the microbeads in solvents, the microbead populations MB1 - MB4 and MB6 were incubated in different solutions (water, acetone, acetone/methanol 1:2, acetone/methanol 1:5, chloroform, chloroform/methanol 1:2, chloroform/methanol 1:5 and methanol) for 1 h at RT and then analysed via VideoScan technology (Rödiger et al., 2012b).

### Coupling of fluorescently labelled phospholipids on hydrophobic microbead surface

Independently, fluorescently labelled phospholipids CL-Cy5, PE-Cy5 and PC-Cy5 were coupled by passive adsorption to the hydrophobic microbead surface of microbead populations MB1-MB5 in 200 μL tubes. Per population, 105 microbeads were washed with 200 μL of 80 % methanol (Carl Roth GmbH + Co.KG, Karlsruhe, Germany) and then incubated at a final concentration of 20 μg/μL (in 80 % methanol) for 3 h at 1,200 rpm and 28 °C in the thermo shaker. This was followed by three washing steps with 1 × tris buffered saline containing 0.01 % Tween 20 (TBS - T) for 1 min each at 13,000 rpm in the centrifuge. For the dilution series, the passive adsorption was performed with different concentrations of fluorescence labelled phospholipids CL-Cy5 and PE-Cy5 (0.1 μg/μL to 100 μg/μL) on microbead populations MB1-MB4.

### Contact angle measurement

For contact angle measurement, coupled and unloaded microbeads were dropped onto a hydrophobic slide (200 μL). These microbeads were dried overnight at RT. Subsequently, one μL of water was added to the dried bead suspension via a contact angle metre and the contact angle was determined by software of OCA 30.

### Digestion of coupled fluorescently labelled phospholipids on the microbeads by specific phospholipases PLA2 and PLC

Microbeads (MB1 - MB5) coupled with fluorescently labelled phospholipids were each incubated with 1 U of a phospholipase (phospholipase A2 in TBS with 20 mM CaCl_2_ and phospholipase C in TBS, Figure 1) for 1 h at 37 °C. The fluorescence signal of the microbeads was measured every 40 sec within 1 h.

**Fig. 1.**
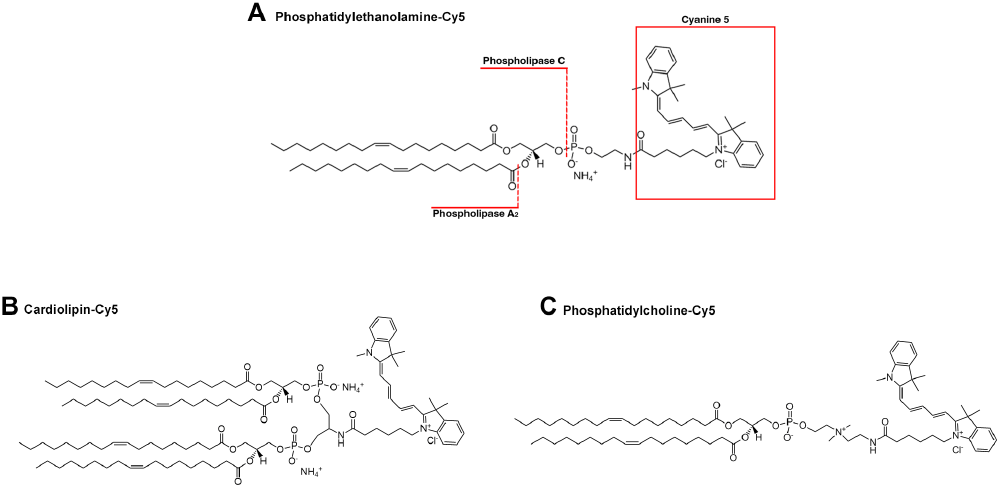
Cutting points of the phospholipases applied. Figure 1 illustrates the structural formulas of the phospholipids labelled with Cy5 PE (A), CL (B), and PC (C). Cutting points of the phospholipases applied (phospholipase A2 and phospholipase C) marked on the example of phosphatidylethanolamine Cy5. Phospholipase A2 cuts the ester bond to the second fatty acid chain at the sn-2 position (Burke and Dennis, 2009; Garcia et al., 2017). Phospholipase C catalyses the hydrolysis of the phosphodiester bond to the glycerol backbone (Garcia et al., 2017).

### Statistical Analysis

The data was analysed with RKWard v. 0.7.5 (Rödiger et al., 2012a). The Tukey HSD was used as statistical tests in which multiple comparisons are taken into account.

## Results and Discussion

The analysis of lipids from human serum is becoming increasingly relevant in diagnostics (Filippas-Ntekouan et al., 2017). In order to quickly diagnose cancer or the antiphospholipid syndrome, it is necessary to have a reaction chamber that is as small as possible, in which little sample material is used, but many parameters can be identified simultaneously (Rödiger et al., 2014). Microbeads are suitable for this purpose. Due to the properties of lipids (Burdge and Calder, 2015; Christie and Han, 2012), these must be stable in solvents and the lipids must be permanently and directionally bound to microbeads. We have developed a way to produce hydrophobic microbeads that are stable in solvents and to couple them successfully in a directed manner and with different phospholipids.

### Solvent stability of carboxylated and hydrophobic microbeads

Lipids are hydrophobic chemical compounds and are thus insoluble in water, but soluble in solvents such as chloroform or methanol (Burdge and Calder, 2015; Christie and Han, 2012). The lipids used (see table 2) were dissolved in chloroform. In order to couple the lipids to the surface of the microbeads, they must be stable in solvents. For this purpose, the microbead populations MB1 - MB4 (hydrophobic) and MB6 (hydrophilic) were incubated in different solutions and subsequently analysed (Fig. 2). Both – hydrophobic and hydrophilic microbeads – are made of PMMA but hydrophobic microbeads are more cross-linked than the hydrophilic microbeads. Conclusively, the outer shell of hydrophobic microbeads is more stable and the microbeads do not swell in solvents. Thus, the hydrophobic microbead populations MB1 - MB4 are stable in all solvents except 100 % chloroform which seems to attack the surface of the microbeads. The hydrophobic microbeads aggregate in both water and chloroform. In water, the aggregation of the microbeads can be explained by the hydrophobic effect. The microbeads have a hydrophobic surface. Thereby, non-polar molecules (hydrophobic microbeads) aggregate in polar solutions (water). We chose methanol as a dilution reagent for the lipids delivered in chloroform for subsequent experiments. In contrast to hydrophobic microbeads, the carboxylated, hydrophilic microbead population MB6 is only stable in water and methanol. However, the fluorescence intensity of the microbead population MB6 decreases over time in methanol. In the process, the microbeads swell and release their fluorescent dyes into the environment. The release of the fluorescent dyes into the surrounding liquid is also clearly visible when the microbeads are incubated in acetone/methanol 1:2 and chloroform/methanol 1:5.

**Table 2.** Biomolecules employed for the development of a phospholipid assay. AP: Avanti Polar Lipids, INC., Dia: Dianova

| Biomolecule | Term | Conjugation | Product number | Supplier |
| --- | --- | --- | --- | --- |
| CL-Cy5 | Cardiolipin-N-(Cyanine 5) | Cy5 | 810388C | AP |
| PE-Cy5 | Phosphoethanolamine-N-(Cyanine 5) | Cy5 | 810335C | AP |
| PC-Cy5 | Phosphocholine-N-(Cyanine 5) | Cy5 | 850483C | AP |

**Fig. 2.**
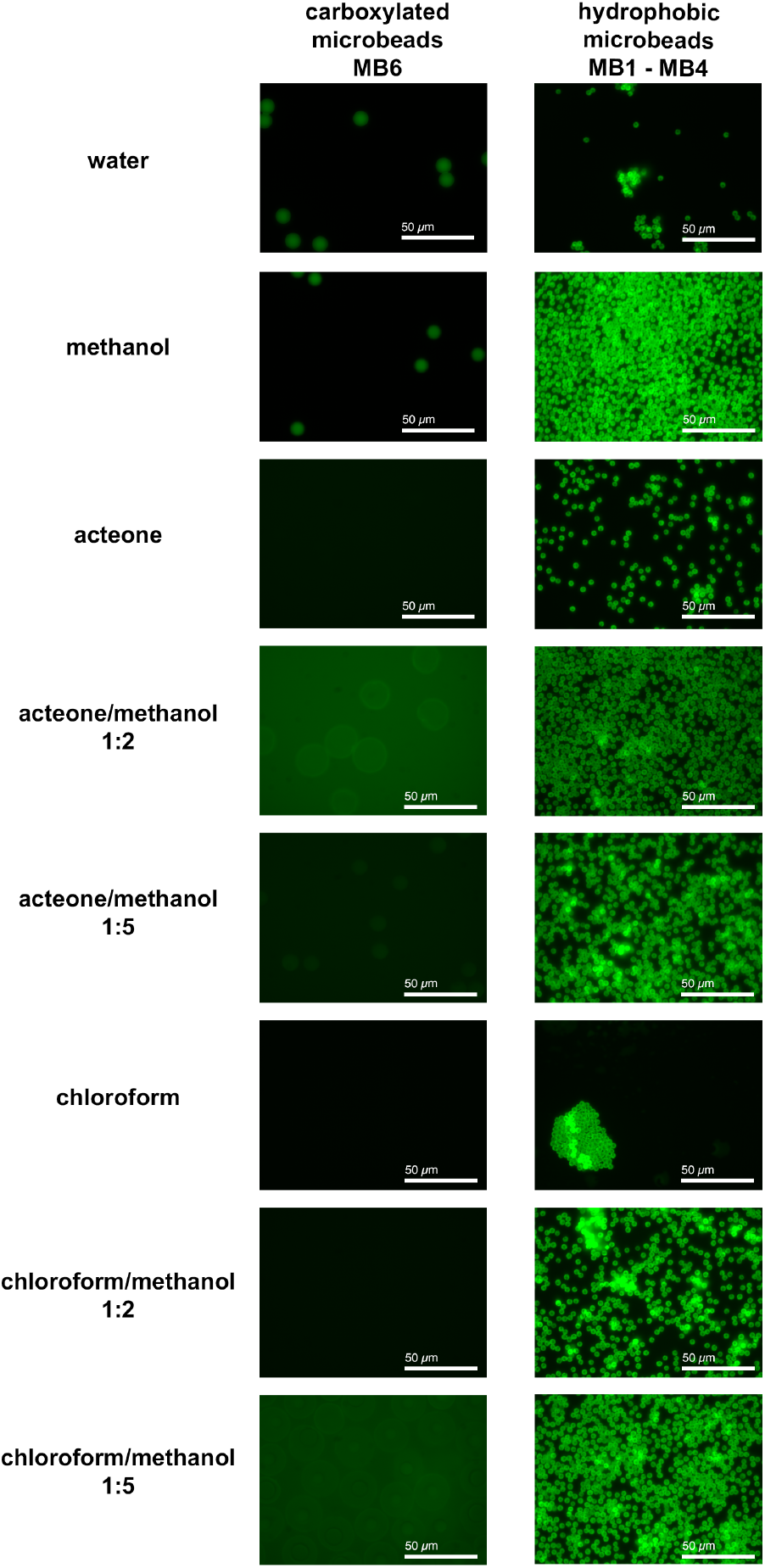
Solvent stability of carboxylated and hydrophobic microbeads. The stability of the microbead populations MB1 - MB4 and MB6 was demonstrated by incubating the microbeads in different solutions (water, methanol, acetone, acetone/ methanol 1:2, acetone/ methanol 1:5, chloroform, chloroform/methanol 1:2 and chloroform/methanol 1:5).

### Coupling of different fluorescently labelled phospholipids to the hydrophobic surface of microbeads

After analysing the solvent stability of the hydrophobic microbeads, the fluorescently labelled phospholipids were coupled to their hydrophobic surface. For this purpose, different fluorescence-labelled phospholipids (CL-Cy5, PE-Cy5 and PC-Cy5; 20 μg/μL) were coupled via passive adsorption to the hydrophobic surface of the microbead populations MB1 - MB4 (Fig. 3). A fluorescence corona around a microbead appeared when phospholipids bound to the surface of the hydrophobic microbeads and the fluorescence signals were quantified by VideoScan technology. The fluorescently labelled phospholipids CL-Cy5, PE-Cy5 and PC-Cy5 bind to the hydrophobic microbead surface and all microbead populations show approximately the same value despite different surface modification (C3 - C18 alkyl-chains). There are only differences in the fluorescence values between the individual phospholipids. A cut-off value of refMFI 0.2 was set for all measurements (calculated from all negative controls). Thus, values above 0.2 are defined as positive. When cardiolipin is bound, the fluorescence value is about 11 times higher than the cut-off value. When PE-Cy5 is coupled, the refMFI value is 16 times the cut-off value and when PC-Cy5 is coupled, the fluorescence value achieved is 12 times the cut-off value. All data is shown in table S1 (Supplementary Note 1). The difference in the fluorescence values of CL-Cy5 at the microbead surface could be explained by the structure. CL-Cy5 differs from the other phospholipids because of the 4 fatty acid chains instead of the usual 2 fatty acid chains of phospholipids (Haines and Dencher, 2002; Houtkooper and Vaz, 2008; Xu et al., 2003). Possibly steric effects occur at the microbead surface and it cannot bind as much CL-Cy5 as PE-Cy5 or PC-Cy5 because of this dimeric nature and cone shape (Houtkooper and Vaz, 2008). The lower fluorescence value of PC-Cy5 to PE-Cy5 could be explained by the behaviour of the PC molecules in 80 % methanol. When the PC molecules interact with water, hydration of the PC occurs and this leads to a sharp decrease in the solubility of the molecule. Hydrated PC molecules tend to aggregate and thus bind more poorly to the microbead surface (Pichot et al., 2013). This experiment shows that different phospholipids can be bound to the hydrophobic surface of the microbeads. The same results could be obtained using flow cytometry as another measurement method (Supplementary Note 1, Fig. S1). Again, the highest fluorescence signal is obtained with the phospholipid phosphatidylethanolamine, followed by phosphatidylcholine and cardiolipin. In the next step, the optimal loading of the fluorescently labelled phospholipids and the associated sensitivity were investigated.

**Fig. 3.**
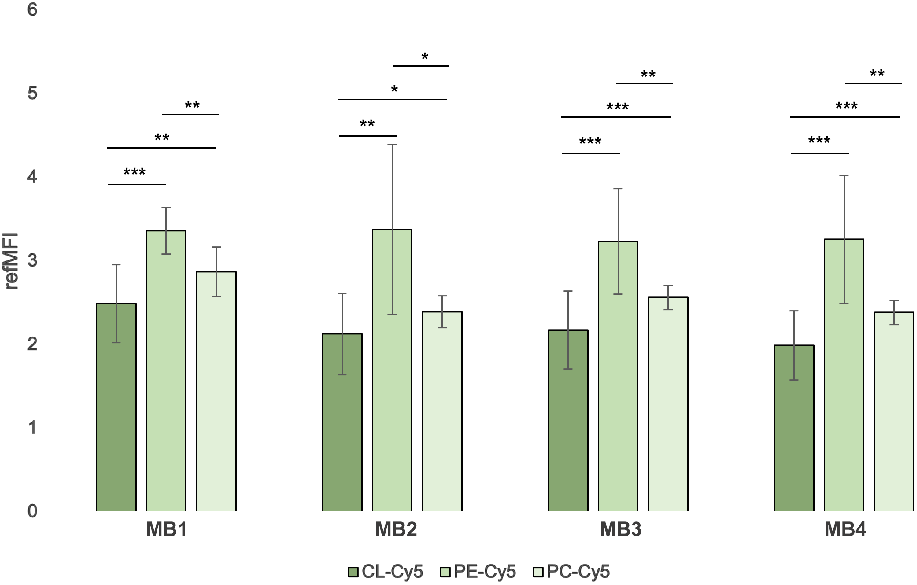
Different fluorescence labelled phospholipids on hydrophobic microbeads. Hydrophobic microbead populations MB1 - MB4 were successfully coupled with 20 μg/μL of phospholipids CL-Cy5, PE-Cy5 and PC-Cy5. The coupling of PE-Cy5 reaches the highest fluorescence signal and CL-Cy5 represents the coupling with the lowest fluorescence signal. All measured signals differ significantly from each other (significances given as * p < 0.05; ** p < 0.01; *** p < 0.001).

### Determination of optimum loading density of CL-Cy5

Optimal loading density of biomolecules on the microbead surface is a prerequisite for the successful development of a detection assay. To be able to define this for further experiments, the phospholipids CL-Cy5, PE-Cy5 and PC-Cy5 were coupled to the microbead surface of MB1 - MB4 in a serial dilution series (0.1 μg/μL to 200 μg/μL) and evaluated using VideoScan technology. As an example, the serial dilution of the fluorescently labelled phospholipid CL-Cy5 (Fig. 4) on hydrophobic microbeads with different surface modifications (C3 - C18 - alkyl chains) is shown here. Based on the fluorescence values obtained, a concentration of 100 μg/μL (MB1 - 20 times, MB2 - MB4 23 times higher than the cut-off) can be defined as the optimal concentration for loading the microbeads. A higher concentration of phospholipid (200 μg/μL) leads to lower fluorescence values due to self-quenching effects of Cy5 (Berlier et al., 2003) or steric effects as well as the Hook effect (Dasgupta and Wahed, 2014; Diaz and Fell, 2004; Pichot et al., 2013) due to the high analyte concentration. A lower concentration of the phospholipid CL-Cy5 results in a signal drop of 52 % and is thus not suitable for optimal microbead loading. An optimal loading density concentration for PE-Cy5 (Supplementary Note 2, Fig. S2) is 20 μg/μL, as the fluorescence values obtained at a higher concentration differ too much (up to 75 %) and thus cannot be used. A minimum detectable fluorescence signal for CL-Cy5 is at a concentration between 2 and 10 μg/μL and for PE-Cy5 at 2 μg/μL.

**Fig. 4.**
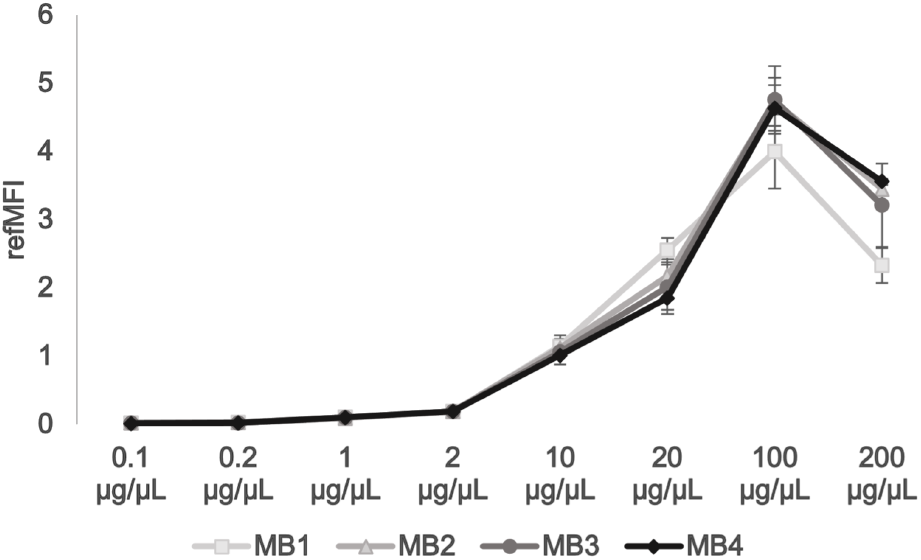
Determination of optimum loading density of CL-Cy5 on hydrophobic microbead populations. Hydrophobic microbead populations MB1, MB2, MB3 and MB4 were coupled with different concentration of CL-Cy5 (0.1 μg/μL to 200 μg/μL) to find out the optimal loading density of the fluorescently labelled phospholipid. The highest fluorescence value is reached at a concentration of 100 μg/μL. From a concentration of 200 μg/μL, the fluorescence values decrease again.

### Verification of directional binding of phospholipids to the microbead surface by phospholipases

It was possible to couple phospholipids to hydrophobic microbeads. In a next step, we tested whether phospholipids bind directionally with the hydrophobic part (the fatty acid residues) to the microbead surface and with the hydrophilic part (head) as freely accessible reaction partner. We chose phospholipases A2 (PLA2) and C (PLC) which can remove the fluorescently labelled head by hydrolysis resulting in a decrease of the fluorescence intensity. PLC catalyses the hydrolysis of the phos-phodiester bond with the glycerol backbone (Garcia et al., 2017). Phospholipase A2 (PLA2) cuts the ester bond at the second fatty acid chain of the phospholipid (Burke and Dennis, 2009; Garcia et al., 2017) (Figure 1). PLA2 is used to show whether the binding of the phospholipids via one fatty acid chain (PE-Cy5 and PC-Cy5) or via two fatty acid chains (CL-Cy5) is sufficient to bind them stably to the microbead surface and thus the fluorescence signal is not reduced (Fig. 5). As shown in Figure 5, addition of PLA2 does not reduce the fluorescence intensity of microbeads, except a very slight decrease due to bleaching effects. Bleaching effects were also visible when microbeads were incubated without enzymes. That incubation with PLA2 does not reduce fluorescence intensity means that binding via only one instead of two fatty acid chains is possible for PE-Cy5 and PC-Cy5 and that binding via only two instead of four fatty acid chains is possible for CL-Cy5. However, as soon as the microbeads are incubated with phospholipase C, the fluorescence signal is reduced. The different timing of the fluorescence decrease may be related to the accessibility of the enzyme or the temperature distribution in the well (37°C - optimal temperature for the enzymes). A device for tempering plates on VideoScan technology was used to perform the experiment. Here, the entire plate is heated and not each individual well, which can lead to temperature fluctuations in individual wells. After about 2000 sec, the fluorescence signal in all populations is significantly reduced compared to the initial signal (Supplementary Note 3, Fig. S3, S4, Table S2). Summarising, the fluorescently labelled phospholipids bind directionally with the hydrophobic part to the hydrophobic microbead surface. The oriented binding of the amphiphilic phospholipids can be explained by two effects: a) the good solvation of the polar hydrophilic domain together with b) hydrophobic interaction (Rego and Patel, 2022) of the lipophilic part with the hydrophobic microbead surface.

**Fig. 5.**
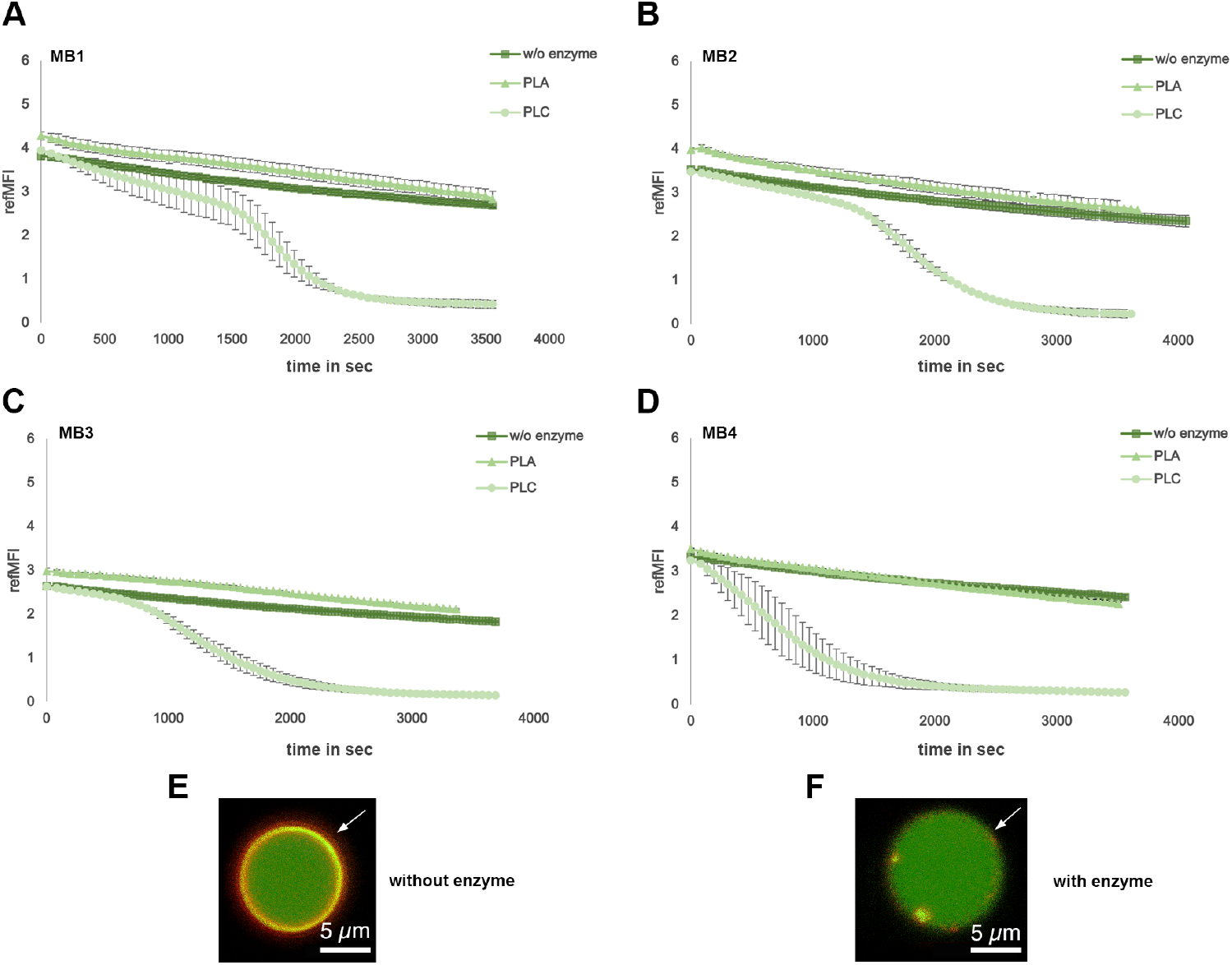
Verification of directional binding of phospholipids on microbead surfaces. Verification of the directional binding of phospholipids using the example of the fluorescently labelled phospholipid PC-Cy5 to the hydrophobic surface of the microbead populations MB1 (A), MB2 (B), MB3 (C) and MB4 (D) by the phospholipases A2 and C. Visible is the initial fluorescence and the associated Cy5 corona on the microbead (E) and the decrease in fluorescence due to hydrolysis and the associated cleavage of the fluorescently labelled head (F). The images were taken using a confocal microscope.

### Broad verification of surface properties via contact angle measurement

The contact angle can be used to determine how hydrophobic a surface is (Law, 2014). Here, the angle from three phases is determined, water droplet, solid surface and air (Kwok and Neumann, 1999). The hydrophobicity of unloaded and coupled microbeads were determined via the contact angle. According to Young’s equation a surface is hydrophobic above or around an angle of 90° and hydrophilic below an angle of 90° (Law, 2014). In comparison, Chen et al., Si et al. and Wang et al. classified hydrophobic surfaces differently. A surface with a contact angle below 10° is superhydrophilic, between 10-65° hydrophilic, between 65-150° hydrophobic and above 150° superhydrophobic (Chen et al., 2022; Si et al., 2018; Wang et al., 2018). We determined the contact angle of uncoated and phospholipid-coated microbeads, respectively. Unloaded microbeads show a contact angle of > 80°, and thus hydrophobicity was confirmed. When phospholipids – labelled (Table 3) and unlabelled (data not shown) - were directionally bound, microbeads were more hydrophilic and all contact angles were between 60 and 83°. According to the contact angle, the phospholipids seem to bind best to MB3, here the contact angle is lowest after coupling. According to the definition of Chen et al., Si et al. and Wang et al. (Chen et al., 2022; Si et al., 2018; Wang et al., 2018, only the surface of microbead population MB3 is hydrophilic, all other microbead surfaces are still in the hydrophobic range. However, a smaller contact angle (difference of up to 31°) occurred after the coupling of phospholipids. The smallest difference from the initial contact angle occurred when CL-Cy5 bound to MB2, whereas the largest difference occurred when PE-Cy5 bound to MB3. Which indicates that the binding of phospholipids changes the surface properties on the microbeads, giving them a more hydrophilic surface. The determination of the contact angle is subject to various conditions, including the surface condition (rough surface, smooth surface). In this context, the contact angles between rough surfaces can differ greatly from smooth surfaces. In the case of microbeads, no smooth surface can be formed due to their spherical shape. In our particular case, despite the microbeads being in one layer, close to each other, a rough surface is formed. Since the microbeads used have different size ratios (5.5 - 12.5 μm).

**Table 3.** Results of contact angle measurement.

| Microbead | uncoated | CL-Cy5 | PE-Cy5 | PC-Cy5 |
| --- | --- | --- | --- | --- |
| MB1 | 94.1° | 75.1° | 74.6° | 70.5° |
| MB2 | 83.1° | 78.6° | 72.0° | 70.3° |
| MB3 | 91.9° | 65.8° | 60.5° | 64.1° |
| MB4 | 104.5° | 82.5° | 75.8° | 79.7° |

### A hydrophobic surface of the microbeads is mandatory to bind phospholipids directionally

The directional binding of phospholipids to hydrophobic microbeads is essential for high quality of assays detecting antiphospholipid antibodies in human patient sera. To confirm directional binding and thus the necessity of hydrophobic microbeads, we coupled CL-Cy5, PE-Cy5 and PC-Cy5 to hydrophilic microbeads with poly-OH group surfaces (Fig. 6). All three phospholipids bound to hydrophilic microbeads and fluorescence intensities were 22-24 times higher than the cut-off value. In contrast to hydrophobic microbeads, PE-Cy5 did not better bind to hydrophilic microbeads than CL-Cy5 and PC-Cy5 (compare Figure 3 and Figure 6). This suggests that phospholipids are rather bound with their hydrophilic heads including the Cy5 label to the hydrophilic microbead surface resulting in a very similar fluorescence intensity. That the hydrophilic head of phospholipids bound to the hydrophilic microbead surface was confirmed using PLC. PLA2 was excluded since PLA2 cuts fatty acid chains but remaining one fatty acid chain (PE-Cy5 and PC-Cy5) or two fatty acid chains (CL-Cy5) were already sufficient to bind these phospholipids to hydrophobic microbeads (see Figure 5). Though incubation of phospholipid-loaded hydrophilic microbeads with PLC, fluorescence intensities did not decrease (Figure 7 A and Supplementary Note 4, Table S3) indicating on the one hand that PLC hydrolysed the phosphodiester bond at the glycerol backbone, but the head with the Cy5 was still bound to the hydrophilic surface of the microbeads or PLC did not cut the phospholipids. Conclusively, a hydrophobic microbead surface is essential to directionally bind phospholipids to microbeads.

**Fig. 6.**
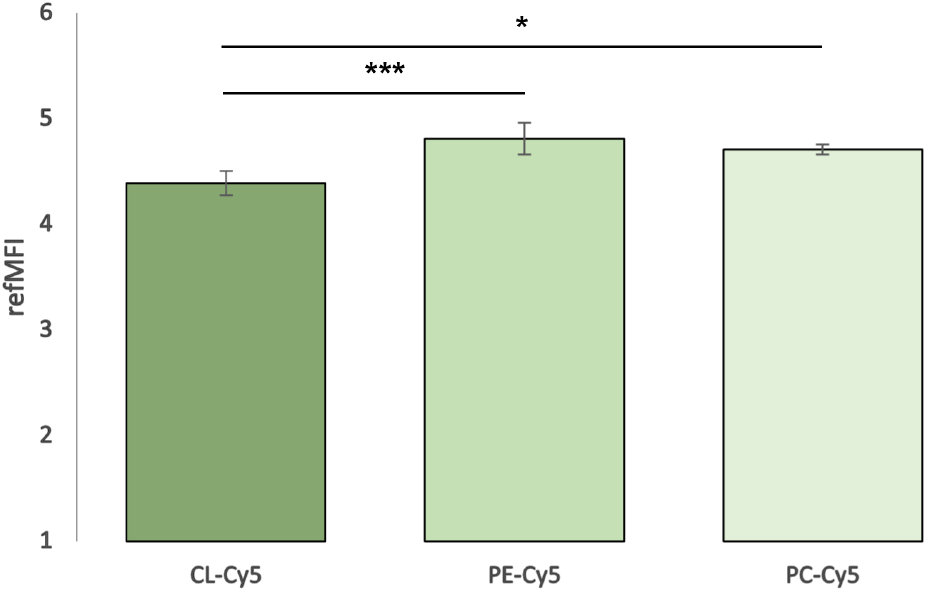
Coupling of hydrophilic microbeads with fluorescently labelled phospholipids. Fluorescently labelled phospholipids CL-Cy5, PE-Cy5 and PC-Cy5 are bound to the surface of hydrophilic microbead populations (MB5). When coupling the phospholipids to hydrophilic microbeads PE-Cy5 shows the highest ligand fluorescence, whereas CL-Cy5 shows the lowest ligand fluorescence upon coupling (significances given as * p < 0.05; ** p < 0.01; *** p < 0.001).

**Fig. 7.**
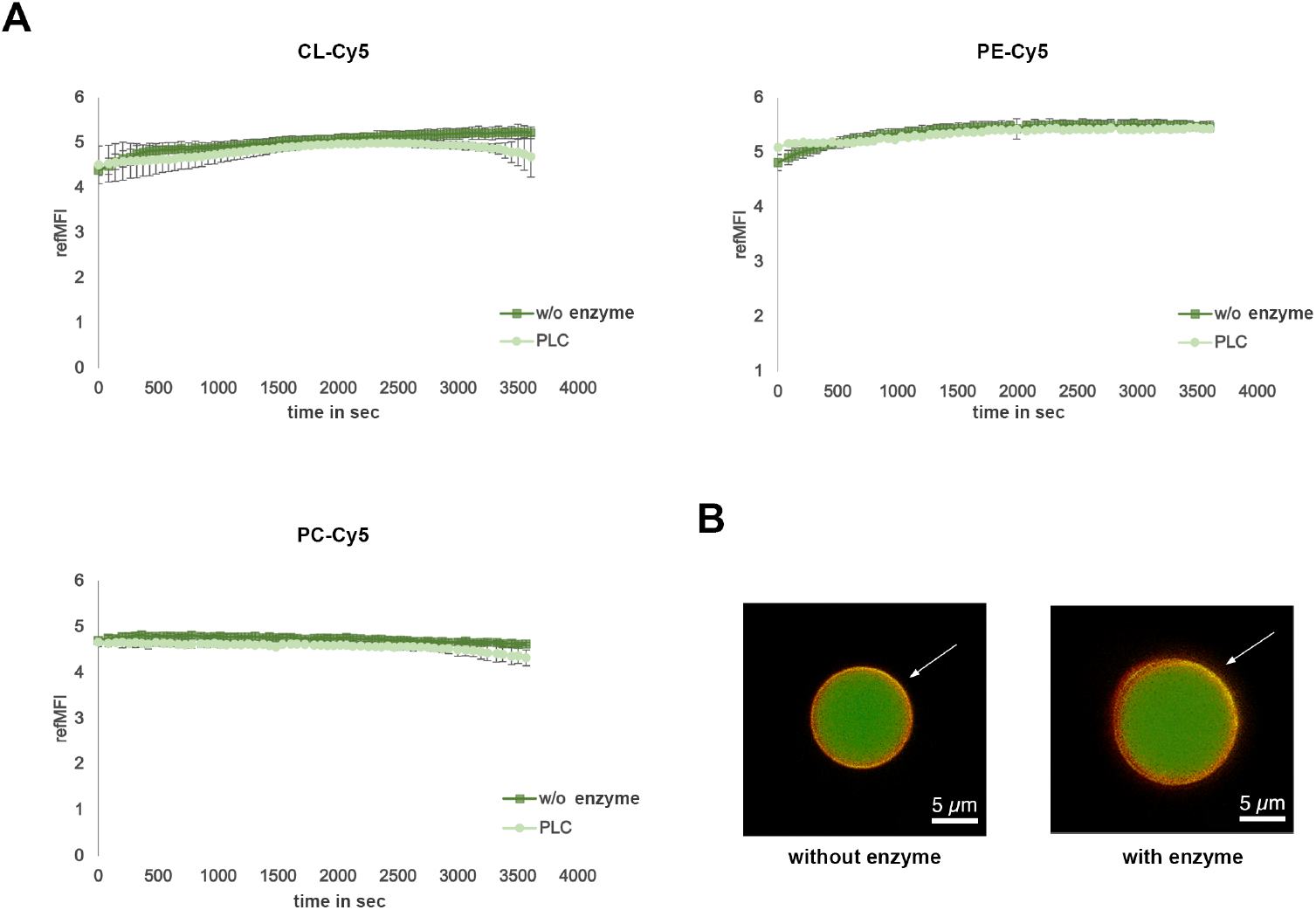
Verification of the necessity of the hydrophobic surface, for the directional binding of the fluorescently labelled phospholipids. Hydrophilic microbeads coupled with fluorescently labelled phospholipids CL-Cy5, PE-Cy5, PC-Cy5 (A) were incubated with phospholipase PLC to show that fluorescence is not reduced at the microbeads and thus it is mandatory to use a hydrophobic microbead surface to bind the fluorescently labelled phospholipids directionally, with the hydrophobic part. The images in 7 B without enzyme and with enzyme also show no decrease in fluorescence. The images were taken using a confocal microscope.

## Conclusions and Outlook

Novel, solvent stable, hydrophobic microbeads were successfully developed and implemented. The microbeads are stable in various solvents and solvent mixtures, except for 100 % chloroform. Fluorescently labelled phospholipids can be directionally and stably coupled to the hydrophobic surface of the microbeads, where the hydrophobic part of the amphiphilic phospholipid is oriented towards the hydrophobic surface of the microbead. However, for this directional binding, a hydrophobic surface of the microbeads is absolutely necessary, as shown by the hydrolysis of the phospholipids by phospholipase C. Contact angle measurements indicate the difference between loaded and unloaded microbeads. Once the phospholipids are bound to the surface of the microbeads, they acquire a more hydrophilic character. Conclusively, a hydrophobic microbead surface is essential to directionally bind phospholipids to microbeads. Directed, immunogenic binding of phospholipids to the microbead surface is a prerequisite for the development of an assay for the detection of antiphospholipid antibodies from human serum. As mentioned at the beginning, these play a major role in antiphospholipid syndrome (APS), among other diseases. We developed a proof-of-principle for the detection of antiphospholipid antibodies against cardiolipin, phosphatidylserine, phosphatidylethanolamine, phosphatidylcholine and phosphatidylinositol. All human sera used were pre-screened by DotBlot (data in Supplementary Note 5, Fig. S6). Furthermore, first experiments regarding the detection of anticardiolipin antibodies on the microbeads could be performed (Supplementary Note 5, Fig. S5). The integration of further phospholipids and human sera has to be performed. The development of such a test would accelerate rapid and accurate diagnosis and save precious and rare patient material from APS patients (incidence of 0.3 % to 1 % of the population) (Cervera et al., 2002; Duarte-Garcia et al., 2019).

## ACKNOWLEDGEMENTS

None.

## COMPLIANCE WITH ETHICAL STANDARDS

The studies on blood material (serum) have been granted by the ethics committee of the Brandenburg University of Technology (BTU) Cottbus-Senftenberg, Cottbus, Germany, (Ethikkommissionssatzung BTU, document number EK2018—3) and were conducted in accordance with the Helsinki Declaration of 1964 (revised 2008). The blood donor gave a written informed consent.

## References

Ahmadpour, S.T., Mahéo, K., Servais, S., Brisson, L., Dumas, J.-F., 2020. Cardiolipin, the Mitochondrial Signature Lipid: Implication in Cancer. Int. J. Mol. Sci. 21, 8031. https://doi.org/10.3390/ijms21218031

Berlier, J.E., Rothe, A., Buller, G., Bradford, J., Gray, D.R., Filanoski, B.J., Telford, W.G., Yue, S., Liu, J., Cheung, C.-Y., Chang, W., Hirsch, J.D., Beechem Rosaria P. Haugland, J.M., Haugland, R.P., 2003. Quantitative Comparison of Long-wavelength Alexa Fluor Dyes to Cy Dyes: Fluorescence of the Dyes and Their Bioconjugates. J. Histochem. Cytochem. 51, 1699–1712. https://doi.org/10.1177/002215540305101214

Burdge, G.C., Calder, P.C., 2015. Introduction to Fatty Acids and Lipids, in: Calder, P.C., Waitzberg, D.L., Koletzko, B. (Eds.), World Review of Nutrition and Dietetics. S. Karger AG, pp. 1–16. https://doi.org/10.1159/000365423

Burke, J.E., Dennis, E.A., 2009. Phospholipase A2 Biochemistry. Cardiovasc. Drugs Ther. 23, 49–59. https://doi.org/10.1007/s10557-008-6132-9

Cervera, R., 2017. Antiphospholipid syndrome. Thromb. Res. 151, S43–S47. https://doi.org/10.1016/S0049-3848(17)30066-X

Cervera, R., Piette, J.-C., Font, J., Khamashta, M.A., Shoenfeld, Y., Camps, M.T., Jacobsen, S., Lakos, G., Tincani, A., Kontopoulou-Griva, I., Galeazzi, M., Meroni, P.L., Derksen, R.H.W.M., de Groot, P.G., Gromnica-Ihle, E., Baleva, M., Mosca, M., Bombardieri, S., Houssiau, F., Gris, J.-C., Quéré, I., Hachulla, E., Vasconcelos, C., Roch, B., Fernández-Nebro, A., Boffa, M.-C., Hughes, G.R.V., Ingelmo, M., 2002. Antiphospholipid syndrome: Clinical and immunologic manifestations and patterns of disease expression in a cohort of 1,000 patients: Clinical and Immunologic Manifestations of APS. Arthritis Rheum. 46, 1019–1027. https://doi.org/10.1002/art.10187

Chen, H., Li, X., Li, D., 2022. Superhydrophilic–superhydrophobic patterned surfaces: From simplified fabrication to emerging applications. Nanotechnol. Precis. Eng. 5, 035002. https://doi.org/10.1063/10.0013222

Christie, W.W., Han, X., 2012. Lipids: their structures and occurrence, in: Lipid Analysis. Elsevier, pp. 3–19. https://doi.org/10.1533/9780857097866.3

Das, U.N., 2020. Can Bioactive Lipids Inactivate Coronavirus (COVID-19)? Arch. Med. Res. 51, 282–286. https://doi.org/10.1016/j.arcmed.2020.03.004

Dasgupta, A., Wahed, A., 2014. Immunoassay Platform and Designs, in: Clinical Chemistry, Immunology and Laboratory Quality Control. Elsevier, pp. 19–34. https://doi.org/10.1016/B978-0-12-407821-5.00002-4

Diaz, M.R., Fell, J.W., 2004. High-Throughput Detection of Pathogenic Yeasts of the Genus Trichosporon. J. Clin. Microbiol. 42, 3696–3706. https://doi.org/10.1128/JCM.42.8.3696-3706.2004

Dimitrijevs, P., Arsenyan, P., 2021. Cardiolipin in the spotlight: Quantitative analysis and fluorescence-based competitive binding assay. Sens. Actuators B Chem. 346, 130537. https://doi.org/10.1016/j.snb.2021.130537

Dinter, F., Burdukiewicz, M., Schierack, P., Lehmann, W., Nestler, J., Dame, G., Rödiger, S., 2019. Simultaneous detection and quantification of DNA and protein biomarkers in spectrum of cardiovascular diseases in a microfluidic microbead chip. Anal. Bioanal. Chem. 411, 7725–7735. https://doi.org/10.1007/s00216-019-02199-x

Dominiczak, M.H., Caslake, M.J., 2011. Apolipoproteins: metabolic role and clinical biochemistry applications. Ann. Clin. Biochem. Int. J. Lab. Med. 48, 498–515. https://doi.org/10.1258/acb.2011.011111

Duarte-García, A, Pham, MM, Crowson, CS, et al. (2019) The Epidemiology of Antiphospholipid Syndrome: A Population-Based Study. Arthritis Rheumatol, 71: 1545–1552.

Falabella, M., Vernon, H.J., Hanna, M.G., Claypool, S.M., Pitceathly, R.D.S., 2021. Cardiolipin, Mitochondria, and Neurological Disease. Trends Endocrinol. Metab. 32, 224–237. https://doi.org/10.1016/j.tem.2021.01.006

Faricelli, R., Esposito, S., Toniato, E., Flacco, M., Conti, P., Martinotti, S., Robuffo, I., 2008. A New Diagnostic Approach to Better Identify Antiphospholipid Syndrome. Int. J. Immunopathol. Pharmacol. 21, 387–392. https://doi.org/10.1177/039463200802100217

Fernandes, A.M.A.P., Messias, M.C.F., Duarte, G.H.B., de Santis, G.K.D., Mecatti, G.C., Porcari, A.M., Murgu, M., Simionato, A.V.C., Rocha, T., Martinez, C.A.R., Carvalho, P.O., 2020. Plasma Lipid Profile Reveals Plasmalogens as Potential Biomarkers for Colon Cancer Screening. Metabolites 10, 262. https://doi.org/10.3390/metabo10060262

Filippas-Ntekouan, S., Liberopoulos, E., Elisaf, M., 2017. Lipid testing in infectious diseases: possible role in diagnosis and prognosis. Infection 45, 575–588. https://doi.org/10.1007/s15010-017-1022-3

Galli, M., Luciani, D., Bertolini, G., Barbui, T., 2003. Anti–β2-glycoprotein I, antiprothrombin antibodies, and the risk of thrombosis in the antiphospholipid syndrome. Blood 102, 2717–2723. https://doi.org/10.1182/blood-2002-11-3334

Garcia, H.S., López-Hernandez, A., Hill, C.G., 2017. Enzyme Technology – Dairy Industry Applications, in: Comprehensive Biotechnology. Elsevier, pp. 608–617. https://doi.org/10.1016/B978-0-12-809633-8.09232-3

Giannakopoulos, B., Passam, F., Ioannou, Y., Krilis, S.A., 2009. How we diagnose the antiphospholipid syndrome. Blood 113, 985–994. https://doi.org/10.1182/blood-2007-12-129627

Haines, T.H., Dencher, N.A., 2002. Cardiolipin: a proton trap for oxidative phosphorylation. FEBS Lett. 528, 35–39. https://doi.org/10.1016/S0014-5793(02)03292-1

Hidaka, H., Hanyu, N., Sugano, M., Kawasaki, K., Yamauchi, K., Katsuyama, T., 2007. Analysis of Human Serum Lipoprotein Lipid Composition Using MALDI-TOF Mass Spectrometry 9.

Houtkooper, R.H., Vaz, F.M., 2008. Cardiolipin, the heart of mitochondrial metabolism. Cell. Mol. Life Sci. 65, 2493–2506. https://doi.org/10.1007/s00018-008-8030-5

Huang, L., Zhang, Y., Su, E., Liu, Y., Deng, Y., Jin, L., Chen, Z., Li, S., Zhao, Y., He, N., 2020. Eight biomarkers on a novel strip for early diagnosis of acute myocardial infarction. Nanoscale Adv. 2, 1138–1143. https://doi.org/10.1039/C9NA00644C

Jizzini, M., Shah, M., Zhou, K., 2020. SARS-CoV-2 and Anti-Cardiolipin Antibodies. Clin. Med. Insights Case Rep. 13, 117954762098038. https://doi.org/10.1177/1179547620980381

Kuromori, Y., Okada, T., Miyashita, M., Harada, K., 2006. Determination of Lipid Composition of Plasma Lipoproteins in Children with a Rapid Agarose Gel Electrophoresis Method. J. Atheroscler. Thromb. 13, 227–230. https://doi.org/10.5551/jat.13.227

Kwok, D.Y., Neumann, A.W., 1999. Contact angle measurement and contact angle interpretation. Adv. Colloid Interface Sci. 81, 167–249. https://doi.org/10.1016/S0001-8686(98)00087-6

Lange, M., Ni, Z., Criscuolo, A., Fedorova, M., 2019. Liquid Chromatography Techniques in Lipidomics Research. Chromatographia 82, 77–100. https://doi.org/10.1007/s10337-018-3656-4

Law, K.-Y., 2014. Definitions for Hydrophilicity, Hydrophobicity, and Superhydrophobicity: Getting the Basics Right. J. Phys. Chem. Lett. 5, 686–688. https://doi.org/10.1021/jz402762h

Loizou, S., McCREA, J.D., Rudge, A.C., Reynolds, R., 1985. Measurement of anticardiolipin antibodies by an enzyme-linked immunosorbent assay (ELISA): standardization and quantitation of results. Clin. exp. Immunol. 62, 738–745.

Lu, S., Yu, T., Wang, Y., Liang, L., Chen, Y., Xu, F., Wang, S., 2017. Nanomaterial-based biosensors for measurement of lipids and lipoproteins towards point-of-care of cardiovascular disease. The Analyst 142, 3309–3321. https://doi.org/10.1039/C7AN00847C

Pichot, R., Watson, R., Norton, I., 2013. Phospholipids at the Interface: Current Trends and Challenges. Int. J. Mol. Sci. 14, 11767–11794. https://doi.org/10.3390/ijms140611767

Plüddemann, A., Thompson, M., Price, C.P., Wolstenholme, J., Heneghan, C., 2012. Point-of-care testing for the analysis of lipid panels: primary care diagnostic technology update. Br. J. Gen. Pract. 62, e224–e226. https://doi.org/10.3399/bjgp12X630241

Postle, A.D., 2012. Lipidomics: Curr. Opin. Clin. Nutr. Metab. Care 1. https://doi.org/10.1097/MCO.0b013e32834fb003

Rego, N.B., Patel, A.J., 2022. Understanding Hydrophobic Effects: Insights from Water Density Fluctuations. Annu. Rev. Condens. Matter Phys. 303–324. https://doi.org/10.1146/annurev-conmatphys-040220-045516

Rödiger, S., Friedrichsmeier, T., Kapat, P., Michalke, M., 2012a. RKWard: A Comprehensive Graphical User Interface and Integrated Development Environment for Statistical Analysis with R. J. Stat. Softw. 49. https://doi.org/10.18637/jss.v049.i09

Rödiger, S., Liebsch, C., Schmidt, C., Lehmann, W., Resch-Genger, U., Schedler, U., Schierack, P., 2014. Nucleic acid detection based on the use of microbeads: a review. Microchim. Acta 181, 1151–1168. https://doi.org/10.1007/s00604-014-1243-4

Rödiger, S., Schierack, P., Böhm, A., Nitschke, J., Berger, I., Frömmel, U., Schmidt, C., Ruhland, M., Schimke, I., Roggenbuck, D., Lehmann, W., Schröder, C., 2012b. A Highly Versatile Microscope Imaging Technology Platform for the Multiplex Real-Time Detection of Biomolecules and Autoimmune Antibodies, in: Seitz, H., Schumacher, S. (Eds.), Molecular Diagnostics, Advances in Biochemical Engineering/Biotechnology. Springer Berlin Heidelberg, Berlin, Heidelberg, pp. 35–74. https://doi.org/10.1007/10-2011-132

Sebastiani, G.D., Iuliano, A., Cantarini, L., Galeazzi, M., 2016. Genetic aspects of the antiphospholipid syndrome: An update. Autoimmun. Rev. 15, 433–439. https://doi.org/10.1016/j.autrev.2016.01.005

Si, Y., Dong, Z., Jiang, L., 2018. Bioinspired Designs of Superhydrophobic and Superhydrophilic Materials. ACS Cent. Sci. 4, 1102–1112. https://doi.org/10.1021/acscentsci.8b00504

Solnica, B., Sygitowicz, G., Sitkiewicz, D., Cybulska, B., Jóźwiak, J., Odrową΀-Sypniewska, G., Banach, M., 2020. 2020 Guidelines of the Polish Society of Laboratory Diagnostics (PSLD) and the Polish Lipid Association (PoLA) on laboratory diagnostics of lipid metabolism disorders. Arch. Med. Sci. 16, 237–252. https://doi.org/10.5114/aoms.2020.93253

Tighe, P.J., Ryder, R.R., Todd, I., Fairclough, L.C., 2015. ELISA in the multiplex era: Potentials and pitfalls. PROTEOMICS – Clin. Appl. 9, 406–422. https://doi.org/10.1002/prca.201400130

Wang, J., Wang, C., Han, X., 2019. Tutorial on lipidomics. Anal. Chim. Acta 1061, 28–41. https://doi.org/10.1016/j.aca.2019.01.043

Wang, X., Hu, L., 2020. Review—Enzymatic Strips for Detection of Serum Total Cholesterol with Point-of-Care Testing (POCT) Devices: Current Status and Future Prospect. J. Electrochem. Soc. 167, 037535. https://doi.org/10.1149/1945-7111/ab64bb

Wang, Y., Ma, K., Xin, J.H.,2018. Stimuli-Responsive Bioinspired Materials for Controllable Liquid Manipulation: Principles, Fabrication, and Applications. Adv. Funct. Mater. 28, 1705128. https://doi.org/10.1002/adfm.201705128

Xu, Y., Kelley, R.I., Blanck, T.J.J., Schlame, M., 2003. Remodeling of Cardiolipin by Phospholipid Transacylation. J. Biol. Chem. 278, 51380–51385. https://doi.org/10.1074/jbc.M307382200

Ye, Y., Chen, Z., Shen, Y., Qin, Y., Wang, H., 2021. Development and validation of a four-lipid metabolism gene signature for diagnosis of pancreatic cancer. FEBS Open Bio 11, 3153–3170. https://doi.org/10.1002/2211-5463.13074

Zou, L., Guo, L., Zhu, C., Lai, Z., Li, Z., Yang, A., 2021. Serum phospholipids are potential biomarkers for the early diagnosis of gastric cancer. Clin. Chim. Acta 519, 276–284. https://doi.org/10.1016/j.cca.2021.05.002

