## Supplement for "Immobilisation of Lipophilic and Amphiphilic Biomarker on Hydrophobic Microbeads"

#### Supplementary Note 1: Coupling of different fluorescently labelled phospholipids to the hydrophobic surface of microbeads

**Table S1: Results of coupling fluorescence labelled phospholipids (in refMFI)**

| Microbeads | CL-Cy5 | PE-Cy5 | PC-Cy5 |
|---|---|---|---|
| MB1 | 2.48 | 3.36 | 2.87 |
| MB2 | 2.12 | 3.37 | 2.39 |
| MB3 | 2.17 | 3.23 | 2.56 |
| MB4 | 1.98 | 3.26 | 2.38 |

#### Flow Cytometry

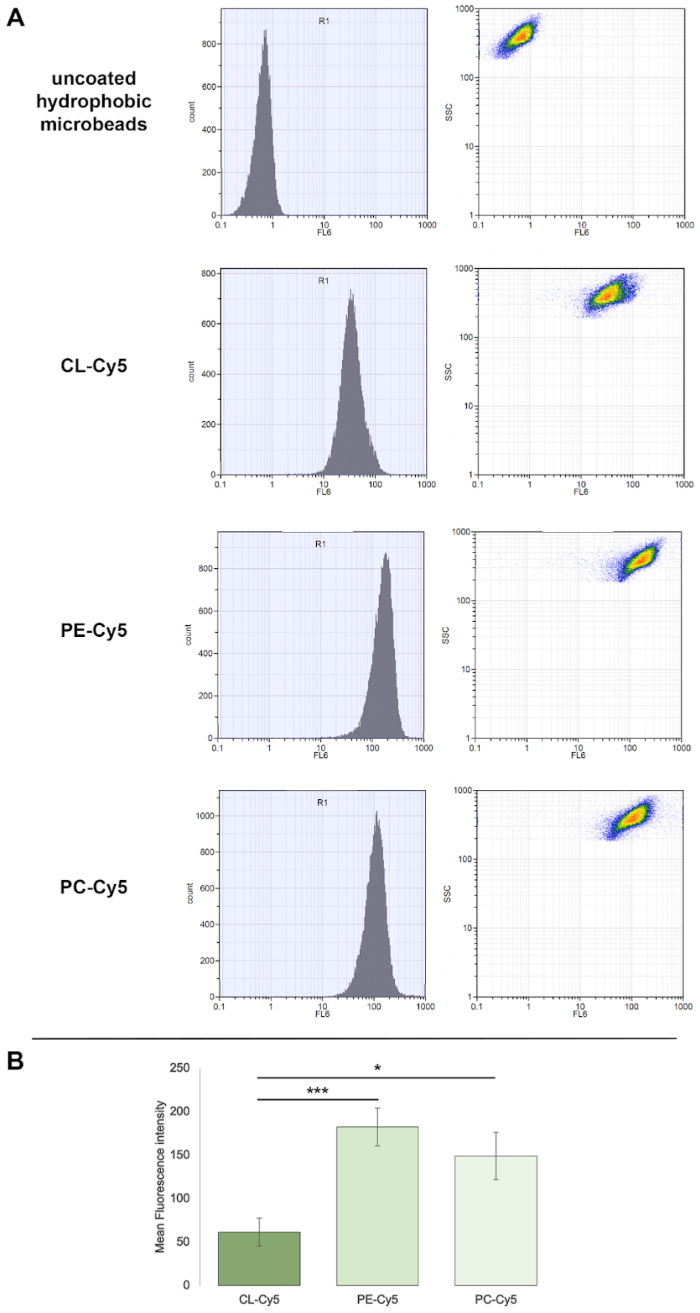


**Fig. S1:**
**Measuring phospholipid coated microbeads with flow cytometry**

Microbeads coupled with fluorescently labelled phospholipids were measured as a control in another system, flow cytometry. Here, a fluorescence signal is also clearly visible on the coupled microbeads in contrast to unloaded microbeads. Compared to the VideoScan technology, the fluorescence values are also lowest for CL-Cy5 and highest for PE-Cy5. Comparable results can be achieved with both technologies (significances given as * $p < 0.05$; ** $p < 0.01$; *** $p < 0.001$).

#### Supplementary Note 2: Optimal Loading Density PE-Cy5

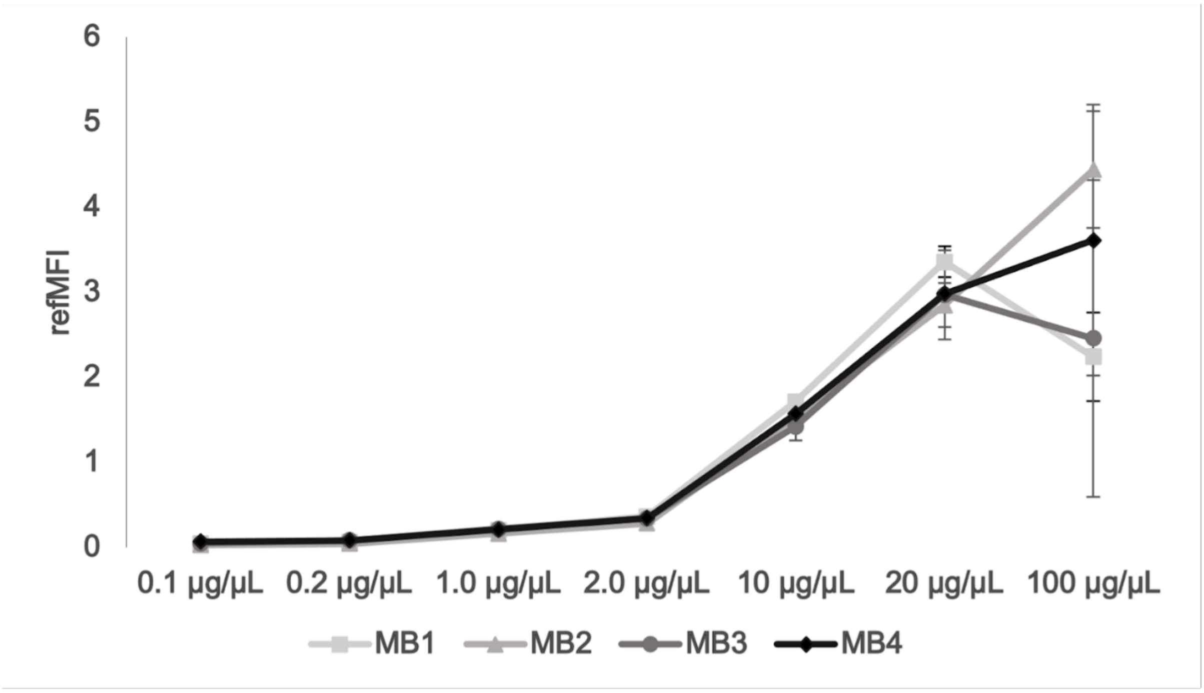


**Fig S2: Determination of optimum loading density of PE-Cy5 on hydrophobic Microbead populations**
Hydrophobic microbead populations MB1 (**A**), MB2 (**B**), MB3 (**C**) MB4 (**D**) were coupled with different concentration of PE-Cy5 (0.1µg/µL to 100 µg/µL) to find out the optimal loading density of the fluorescently labelled phospholipid.

#### Supplementary Note 3: Verification of directional binding of phospholipids to the microbead surface by phospholipases

### CL-Cy5

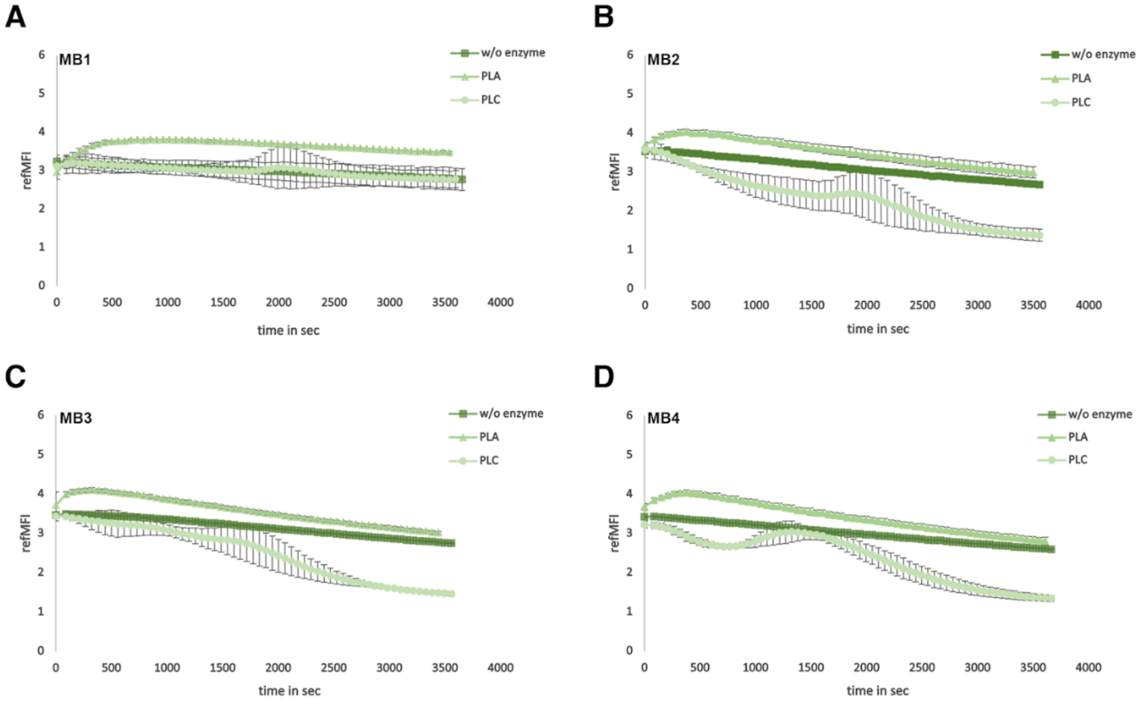


**Fig. S3: Verification of directional binding of phospholipids on microbead surfaces**
Verification of the directional binding of phospholipids using the example of the fluorescently labelled phospholipid CL-Cy5 to the hydrophobic surface of the microbead populations MB1 (**A**), MB2 (**B**), MB3 (**C**) and MB4 (**D**) by the phospholipases A2 and C.

## PE-Cy5

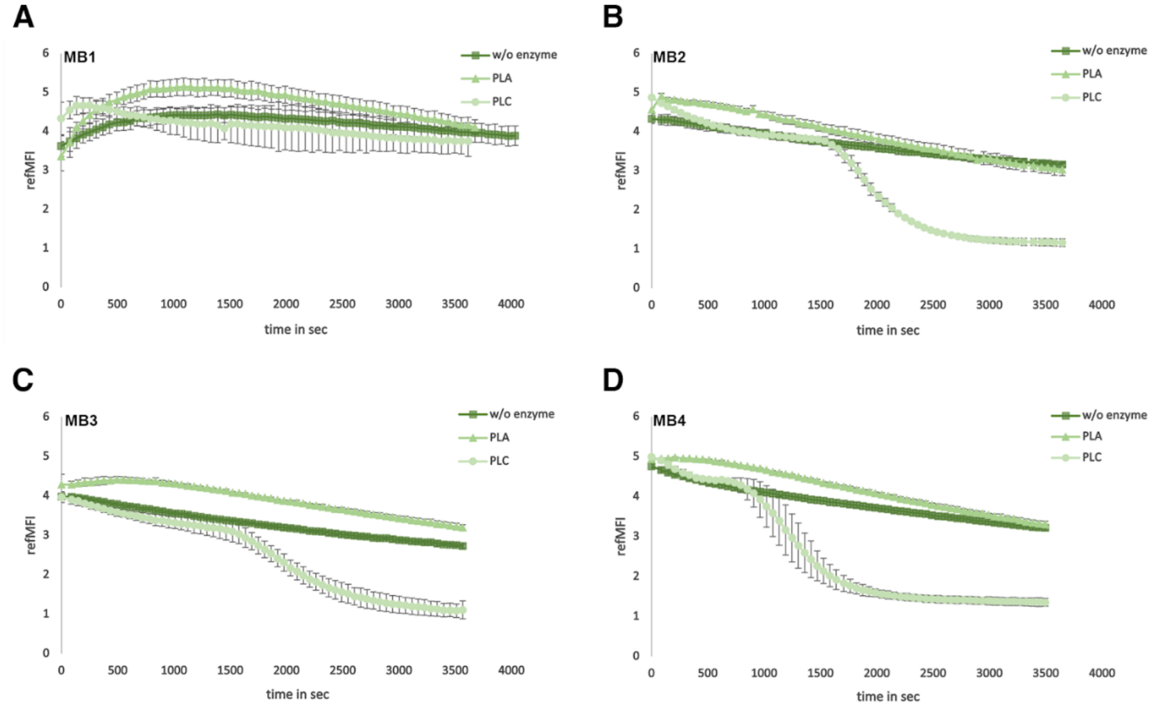


**Fig. S4: Verification of directional binding of phospholipids on microbead surfaces**
Verification of the directional binding of phospholipids using the example of the fluorescently labelled phospholipid PE-Cy5 to the hydrophobic surface of the microbead populations MB1 (**A**), MB2 (**B**), MB3 (**C**) and MB4 (**D**) by the phospholipases A2 and C.

**Table S2: Results of phospholipase C hydrolysis (in refMFI)**

| Microbeads | CL – Cy5 | | PE-Cy5 | | PC-Cy5 | |
|---|---|---|---|---|---|---|
| | before | after | before | after | before | after |
| MB1 | 3.12 | 2.78 | 4.33 | 3.75 | 3.94 | 0.42 |
| MB2 | 3.56 | 1.36 | 4.87 | 1.16 | 3.47 | 0.22 |
| MB3 | 3.43 | 1.42 | 3.97 | 1.11 | 2.62 | 0.14 |
| MB4 | 3.22 | 1.33 | 4.99 | 1.34 | 3.24 | 0.24 |

#### Supplementary Note 4: A hydrophobic surface of the microbeads is mandatory to bind phospholipids directionally

**Table S3: Results of phospholipase C hydrolysis (in refMFI)**

| Microbeads | CL-Cy5 | | PE-Cy5 | | PC-Cy5 | |
|---|---|---|---|---|---|---|
| | before | after | before | after | before | after |
| MB5 | 4.50 | 4.68 | 5.09 | 5.42 | 4.66 | 4.32 |

**Supplementary Note 5: Detection of antiphospholipid antibodies from human serum**

**Method**

### Detection of anti phospholipid antibodies in human sera

Hydrophobic microbeads coupled with unconjugated phospholipids (table S4) were incubated with 1:40 diluted sera (in TBS - T containing 0.01% Tween 20, Table 3) for 1.5 h at 28 °C and 1,200 rpm in the thermo shaker and subsequently washed with TBS - T 3 x for 1 min each at 13,000 rpm in the centrifuge. Subsequently, microbeads were incubated with a fluorescently labelled secondary antibody (anti human IgG - 1:100 - 7.5 µg/µL) for 1 h at 1,200 rpm and 28 °C in the thermo shaker and washed 3 x with TBS - T (0.01 % Tween 20) for 1 min each at 13,000 rpm in the centrifuge. The resulting fluorescence signal was measured using VideoScan technology.

**Table S4: Human sera, their specificity and reactivity**

| Human Serum | Specificity | Reactivity (antiphospholipid antibodies) |
|---|---|---|
| S1 | IgG positive | CL, PS |
| S2 | IgG positive | CL, PS |
| S3 | IgG positive | CL, PS |
| S4 | IgG negative | - |
| S5 | IgG negative | - |
| S6 | IgG negative | - |
| S7 | IgG negative | - |
| S8 | IgG negative | - |

**Results**

### Detection of antiphospholipid antibodies from human serum

In a final step we established a new diagnostic microbead assay for the detection of human antiphospholipid antibodies. The antiphospholipid syndrome is a rare disease with an incidence of 0.3 to 1 % of the population. We assumed that non-conjugated phospholipids behave similarly to fluorescently labelled phospholipids since the fluorophore is located at the head group of the phospholipid and thus should not influence the binding via the hydrophobic fatty acid chains to the microbead surface. We coupled CL to hydrophobic microbeads (populations MB1-MB4). After blocking with 2.5 % BSA TBS-T microbeads were incubated with patient sera positive for anticardiolipin antibodies and with control sera. Positivity and negativity of patient and control sera were determined using a DotBlot (Fig. S6). The bound anticardiolipin antibodies were then detected by an anti-human fluorescently labelled (Alexa-Fluor647) secondary antibody (Fig. S5). The three anticardiolipin antibody-positive sera generated a positive signal on the microbeads with a signal above the cut-off of 0.2. So far, the fluorescence values were low compared to the fluorescence values using the fluorescently labelled phospholipids. This requires optimisation with regard to the use of sera or the coupling of phospholipids. Signal amplification methods such as the use of aptamers or similar could also be suitable. The detection of antiphospholipid antibodies from human serum currently serves as a proof-of-principle and should be further optimised and verified with more human serum and other phospholipids.

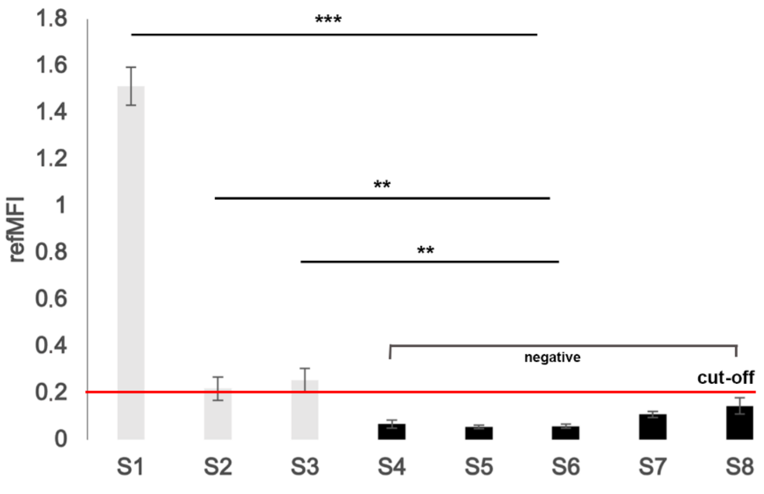


**Fig. S5: Detection of antiphospholipid antibodies in human sera**
Different hydrophobic microbead populations were coupled with phospholipid CL, then incubated with 8 different patient sera (S1 - S8) and the bound antibodies from the serum were detected using a fluorescently labelled anti-human secondary antibody. The human sera S1-S3 were positive (the fluorescence values are above the cut-off (mean + 3*SD, red line)). The fluorescence values of the human sera S4-S8 are below the cut-off and are therefore negative (significances given as * $p < 0.05$; ** $p < 0.01$; *** $p < 0.001$).

#### Dot Blot

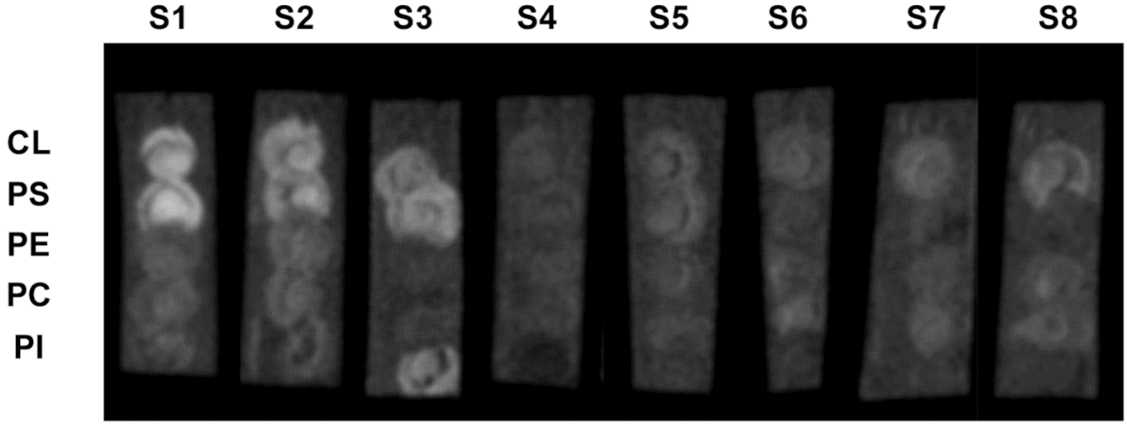


**Fig. S5: Dotblot analysis of phospholipid antibodies in human sera**
On a hybond PVDF membrane 1µL of different unconjugated phospholipids (CL, PS, PE, PC, PI) were dropped and dried for 1h. Subsequently, the membrane was blocked with 2.5% BSA TBS for 30 min. DotBlots were incubated in 1:40 diluted (in 2.5% BSA TBS) human serum (S1-S8) for 1.5 h. After incubation, these were washed 3 x 10 min with TBS-T (0.01% Tween 20) and incubated with a fluorescently labelled secondary antibody against human IgG (1:100 in 2.5 % BSA TBS) for 1 h. Subsequently, the blots were washed 3 times with TBS -T (0.01% Tween 20) and analysed using the Biostep S chemiluminescence imager. Only human sera S1-S3 were positive for antiphospholipid antibodies, all others show negative results.

**Table S1: Analysis of the brightness values in DotBlot using Image J**

| Sera | CL | PS | PE | PC | PI |
|---|---|---|---|---|---|
| S1 | 137.37 | 147.98 | 79.94 | 80.55 | 72.21 |
| S2 | 121.67 | 116.99 | 85.38 | 83.59 | 58.63 |
| S3 | 105.42 | 118.33 | 58.19 | 56.66 | 95.84 |
| S4 | 61.60 | 60.36 | 60.83 | 56.37 | 33.59 |
| S5 | 72.26 | 78.93 | 65.97 | 64.22 | 53.85 |
| S6 | 73.60 | 60.29 | 66.33 | 80.53 | 51.22 |
| S7 | 84.06 | 56.99 | 68.33 | 79.08 | 54.55 |
| S8 | 83.78 | 56.81 | 73.52 | 75.13 | 51.75 |

A threshold of 90 must be reached for a positive signal and thus the detection of phospholipid antibodies in human serum.
